# The gut microbiota of rural and urban individuals is shaped by geography and lifestyle

**DOI:** 10.1101/2020.03.19.999656

**Authors:** Mubanga Hellen Kabwe, Surendra Vikram, Khodani Mulaudzi, Janet K. Jansson, Thulani P. Makhalanyane

## Abstract

Understanding the structure and drivers of gut microbiota remains a major ecological endeavour. Recent studies have shown that several factors including diet, lifestyle and geography may substantially shape the human gut microbiota. However, most of these studies have focused on the more abundant bacterial component and comparatively less is known regarding fungi in the human gut. This knowledge deficit is especially true for rural and urban African populations. Therefore, we assessed the structure and drivers of rural and urban gut mycobiota. Our participants (n=100) were balanced by geography and sex. The mycobiota of these geographically separated cohorts was characterized using amplicon analysis of the Internal Transcribed Spacer (ITS) gene. We further assessed biomarker species specific to rural and urban cohorts. In addition to phyla which have been shown to be ubiquitous constituents of gut microbiota, Pichia were key constituents of the mycobiota. We found that several factors including geographic location and lifestyle factors such as the smoking status were major drivers of gut mycobiota. Linear discriminant and the linear discriminant analysis effect size analysis revealed several distinct urban and rural biomarkers. Together, our analysis reveals distinct community structure in urban and rural South African individuals. Geography and lifestyle related factors were shown to be key drivers of rural and urban gut microbiota.

**Importance:** The past decade has revealed substantial insights regarding the ecological patterns of gut microbiomes. These studies have shown clear differences between the microbiomes of individuals living in urban and rural locations. Yet, in contrast to bacteria we know substantially less regarding the fungal gut microbiota (mycobiome). Here we provide the first insights regarding the mycobiome of individuals from urban and rural locations. We show that these communities are geographically structured. Further we show that lifestyle factors, such as diet and smoking, are strong drivers explaining community variability.

## Introduction

By comparison to prokaryotes (bacteria and archaea), eukaryotes are considered part of the rare “biosphere” of the gut (1, 2). Despite their low abundances, fungi play significant roles in host physiology (2-5). Recent studies have shown that the gut fungal community composition is less stable over time, compared to bacterial communities (4, 6, 7). These studies suggest that the gut mycobiota is more variable than bacterial communities, and may be influenced substantially by environmental factors (3, 7). Despite evidence confirming the gut microbiota is diverse and interacts with the host immune system (3, 8, 9), knowledge regarding the community structure of healthy human gut mycobiota remains scant.

Most studies have focused on the potential roles played by the mycobiota in the aetiology of gut diseases (10-12). These studies have provided crucial insights on the role of the mycobiota as a potential drivers of immunological disorders and as opportunistic pathogens in immunocompromised hosts (13). Further, dysbiosis of gut mycobiota has been linked to obesity, colorectal cancer and Inflammatory Bowel Diseases (IBDs) (12, 14, 15). Decreased abundances of *Saccharomyces cerevisiae* and higher proportions of *Candida albicans* were found in IBD patients compared to healthy controls. A recent study showed that Crohn’s disease-specific gut environments may select for fungi to the detriment of bacteria suggesting disease-specific inter-kingdom network alterations in IBD (12). Yet, despite these beneficial effects, there remains a clear deficit in knowledge regarding the precise role played by the gut mycobiota in disease prevention. Relatedly, the factors which drive the diversity and community structure of gut mycobiome remain underexplored. Assessing the influence of environmental factors on the gut mycobiome across a wider cohort of participants is crucial for determining the effects on host-microbiota dynamics and health.

Several studies have evaluated the effects of age (16-18), gender (17), diet (19), diabetes and obesity (15, 20, 21), anorexia nervosa (22), differences across body sites (23, 24) and geographical locations (6, 25, 26) on mycobiome composition and diversity. Yet, these studies are mostly disease centric or focussed on Asian (26) and/or Western populations (6, 7, 19). To our knowledge, only one study has investigated the gut eukaryotic diversity of African individuals (27). Although these studies improved our understanding of the mycobiome, there may be several confounding factors such as genetic differences. These differences make it difficult to assess, for instance, the effects of living in urban or rural areas on the microbiome. The effects of diet, geographic locality and lifestyle, on the gut microbiome are often assumed but rarely examined. Where these relationships are assessed, the majority of studies have primarily focused on the ecologically abundant bacteria (28, 29) with assertions that their patterns will likely hold for other taxa, including mycobiomes.

Here, we applied amplicon sequencing of the fungal internal transcribed spacer (ITS) of the rRNA gene on samples collected from individuals living in urban and rural areas in Africa. We provide the first insights regarding the drivers of mycobiome community structure and potential biomarkers specific to individuals from urban and rural locations. Previous studies of the gut mycobiome have primarily investigated small cohorts with fewer than 20 individuals (25, 30, 31) with very few studies investigating larger cohorts (6, 32). This study represents the first analysis of the faecal mycobiota in a large cohort of healthy sub-Saharan individuals (100 volunteers). Furthermore, this is the first study which compares the composition and diversity of the gut mycobiome of geographically separated non-western individuals with the same ethnicity. We further explored potential biomarker taxa in urban and rural individuals and explore how these taxa vary between the two areas. Using extensive predictor variables collected from participants, we show that geography and lifestyle structure the gut mycobiome of rural and urban South African individuals.

## Results

### Similarities and differences between urban and rural individuals

We assessed the faecal mycobiota of South African adults living in rural (*n*=50) and urban regions (*n*= 50) by assessing stool samples (see details regarding sample recruitment in Methods). We recruited an equal number of male and female volunteers across two villages in the Limpopo region of South Africa, which is located roughly 500km from the urban site (Pretoria) (**Figure 1a**). To gain insights regarding predictive variables which may shape the gut microbiome, detailed questionnaires were distributed to all volunteers. The volunteers from Ha-Ravele and Tshikombani villages (population size of roughly 200,000, representing the rural participants) were on average 24 years (mean ± 6.3). Volunteers from Pretoria (total population of approximately 2.1 million) were on average 31 years (mean ± 9.1). The BMI of all participants was above 26.45, resulting in a cohort of individuals classified as obese, less than 15.9% of participants were smokers.

**Figure 1.**
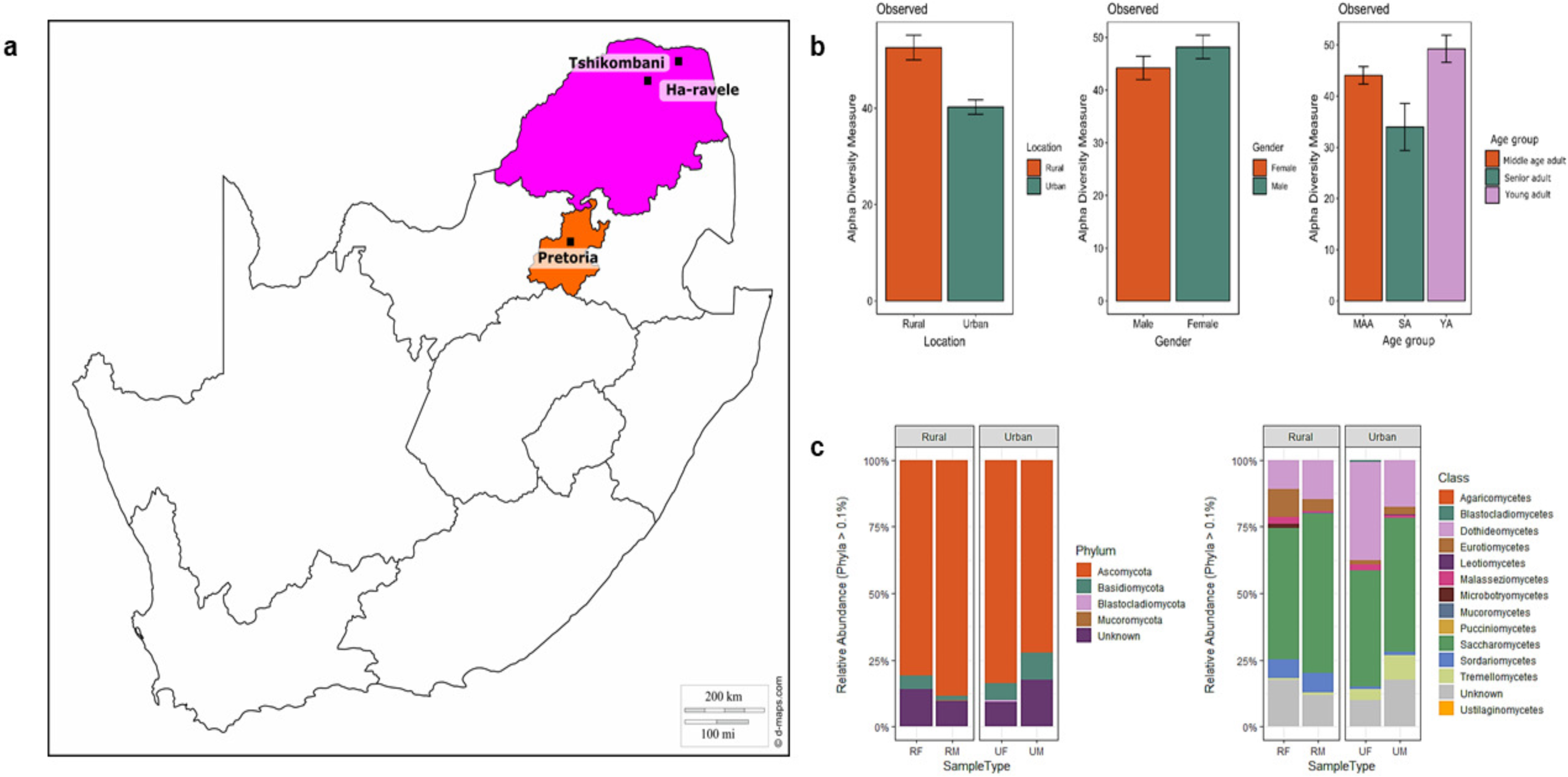
Geographic locations and diversity estimates **a**) The three sampling locations in Gauteng (Pretoria) and Limpopo (Ha-ravele and Tshikombani) provinces of South Africa **b**) The differences in mycobiota species richness between the two locations, gender and age group and, **c**) The relative abundances of taxa at phylum and class levels within each location. The abundance of each taxon was calculated as the percentage of sequences per gender (RF = Rural female, RM = Rural male, UF = urban female and UM = Urban male) from each location for a given microbial group. The group designated as ‘Unknown’ encompasses unclassified sequences together with classes representing > 0.1% of the total sequences. The bar size represents the relative abundance of specific taxa in the particular group, with colours referring to taxa according to the legend.

Amplicon sequence data from 95 volunteers (samples from 5 rural volunteers were excluded due to low quality reads) generated 5,936,454 raw reads. A total of 5,414,023 fungal reads were retained after sequence filtering and chimera removal. Of these, 1,636,180 reads were assigned to OTUs and these were further clustered into 1,911 OTUs using a 97% cut-off. A higher proportion of fungal OTUs were unique to location with urban and rural samples accounting for 47.9% and 45.3% of reads, respectively (**Supplemental Figure 1a**).

After random subsampling to the lowest read count, 155 OTUs were excluded from further analyses. The resulting accumulation curves showed reasonable sequence saturation, at a regional level (**Supplemental Figure 1b**). Fungal species richness was significantly higher in the stool samples of rural volunteers (Observed; W = 615.5, *p-*Value = 1.471e4) compared to urban volunteers (**Figure 1b**). However, we did not find significant differences in richness based on gender and age group.

### Two ubiquitous fungal phyla in urban and rural locations

Overall, four distinct fungal phyla were detected in urban and rural gut mycobiomes, based on sequences with relative abundances above 0.1% (**Figure 1c**). The majority of sequences were assigned to members of the phyla *Ascomycota* and *Basidiomycota*, that constituted 80.7% and 6.1% of the total relative abundance, respectively. Unknown sequences constituted 12.9% of the fungal mycobiome. In total, 13 distinct fungal classes were identified with *Saccharomycetes* constituting the majority of sequences (50.1%) followed by *Dothideomycetes* (20.3%), *Eurotiomycetes* (4.8%) and *Sordariomycetes* (3.9%). Unknown fungal genera dominated our cohort (18.4%), followed by *Pichia* (17.6%) *Candida* (17.1%) and *Cladosporium* (5.9%). However, no significant difference was found between taxa abundance at the class level for the gut mycobiota of rural and urban participants (W = 1054, *p*-value = 0.5992). The difference between the gut mycobiota of rural and urban individuals, across the two locations, was not statistically significant (Kruskal-Wallis chi-squared = 2.9875, df = 3, *p*-value = 0.3936).

To assess the degree of uniqueness of a given sample in relation to the overall community composition, we assessed the local contribution to beta diversity (LCTBD). We found that samples from urban volunteers contributed a greater fraction of the overall community diversity (*p*-Value < 0.05). Samples with high local contribution to beta diversity (LCTBD) had a high abundance of *Basidiomycota* and other unknown taxa. In contrast, only two samples from the rural location contributed to overall community diversity (**Supplemental Figure 1c**).

### Distinct mycobiota among urban and rural volunteers unrelated to gender

Differences in the fungal community structure between the rural and urban localities were visualized in an NMDS plot (**Figure 2a**). Urban and rural samples formed distict clusters [PERMANOVA (R^2^ = 0.070; *p*-Value = 0.0001), ANOSIM (R = 0.43, *p*-Value = 0.001) and ADONIS (R^2^ = 0.07034 *p*-Value = 0.0001)]. However, male and female samples did not cluster separately. Pairwise analysis of permutational multivariate analysis of variance (PERMANOVA) showed that there was no significant difference between female vs male urban, and female vs male rural individuals (R^2^ = 0.018; R^2^ = 0.023; respectively and *p*-Value > 0.4 for both). However, there was a significant difference between the gut mycobiota of urban versus rural female and male participants (R^2^ < 0.074 for all; *p*-Value = 0.001 for all).

**Figure 2.**
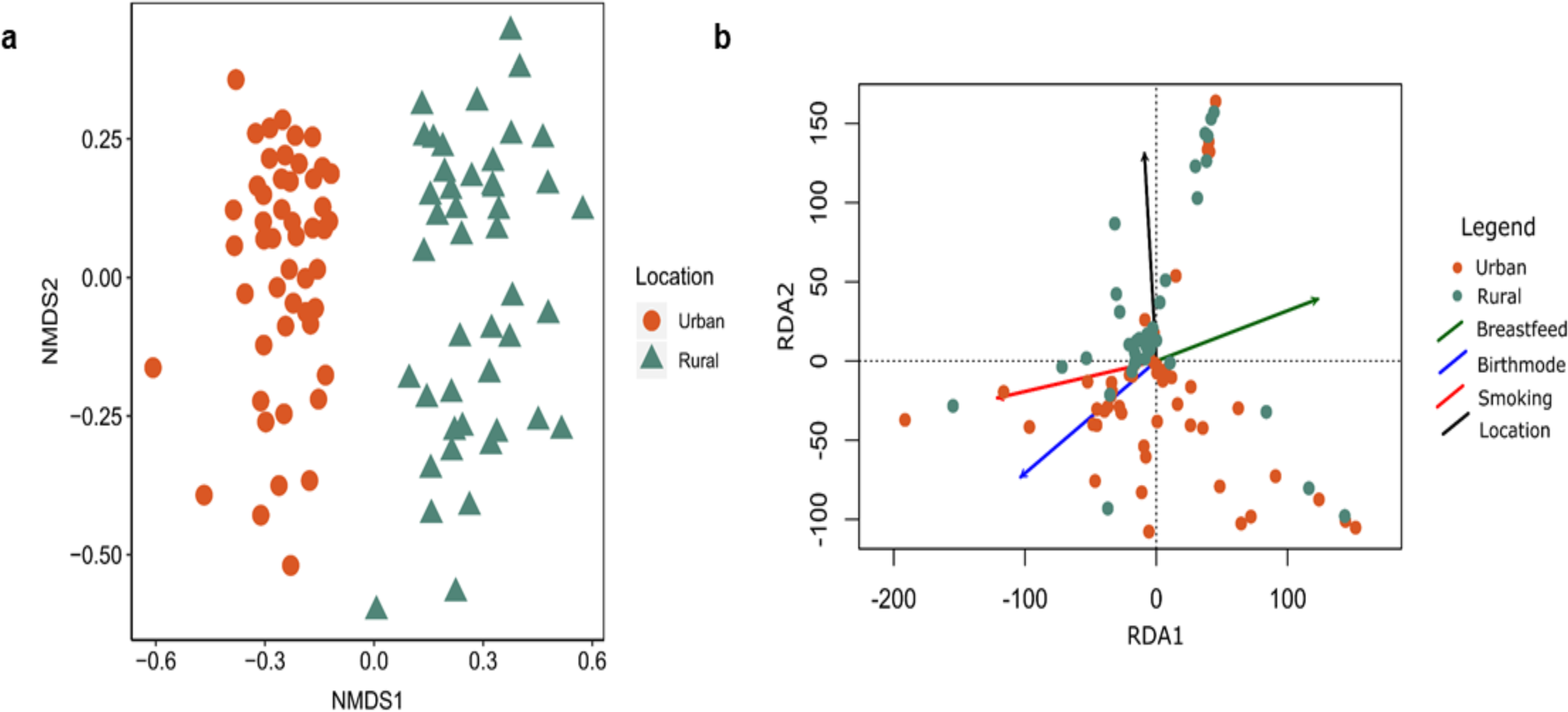
Overview of mycobiota structure and significant environmental drivers **a**) The non-metric multidimensional scaling (NMDS) plot based on Bray–Curtis dissimilarity and, **b**) Redundancy analysis (RDA) showing community structure in response to four selective variables. The filled shapes reflect fungal community composition in the different locations, with colours referring to location and the different explanatory variables according to the legend.

### Ecological drivers of gut mycobiota

Redundancy analysis (RDA) was performed to determine which predictor variables significantly explained the variation in fungal community composition (**Figure 2b**). Four predictive variables were significant (r^2^ > 0.2; *p*-Value < 0.05) drivers of community composition and structure. Predictive variables included; breastfeeding, smoking, mode of birth and location; all of which significantly influenced the fungal community composition.

We conducted correlation analyses to explore the relationships among dominant gut species. Our results showed a few highly positive correlations in the rural cohort: between *Mucor* and *Dipodascus, Mucor* and *Naganishia, Clavispora* and *Lentendrea*, and between *Udeniomyces and Lentendrea* (**Figure 4a**). Whereas, the strongest negative correlation was found between *Dipodascus* with *Trichoderma, Dipodascus* with *Ascotricha* and *Dipodascus* with *Chalastospora*. Within the urban cohort, *Xeromyces* and *Agaricus, Diutina* and *Clavispora*, and *Dekkera* and *Diutina* exhibited the strongest positive correlations (**Figure 4b**). The strongest negative correlations were detected between *Clavispora* with *Filobasidium*, and with *Verrucaria* and *Malassezia*.

**Figure 3.**
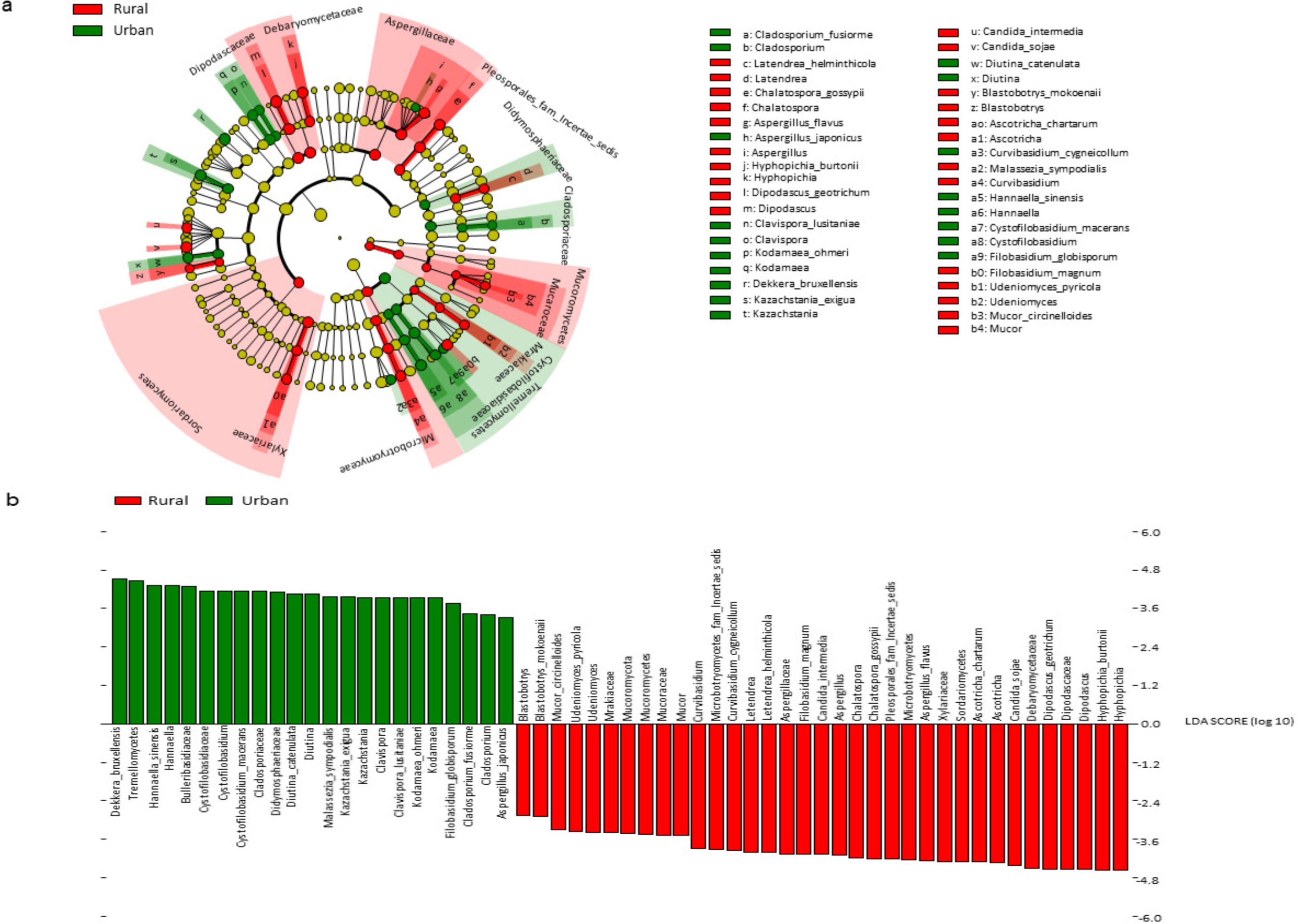
The results of Linear discriminant analysis (LDA) effect size (LefSe) analysis of rural and urban gut mycobiota **a)** The cladogram shows the output of the LEfSe algorithm, which identifies taxonomically consistent differences between rural (Ha-ravele and Tshikombani villages) and urban (Pretoria) fungal community members, respectively. Taxa with nonsignificant differences are represented as yellow circles and the diameter of the circle is proportional to relative abundances **b**) The histogram of the LDA scores was computed for differentially abundant taxa between the rural and urban gut mycobiota. The bar size represents the effect of the size of specific taxa in the particular group at species level

**Figure 4.**
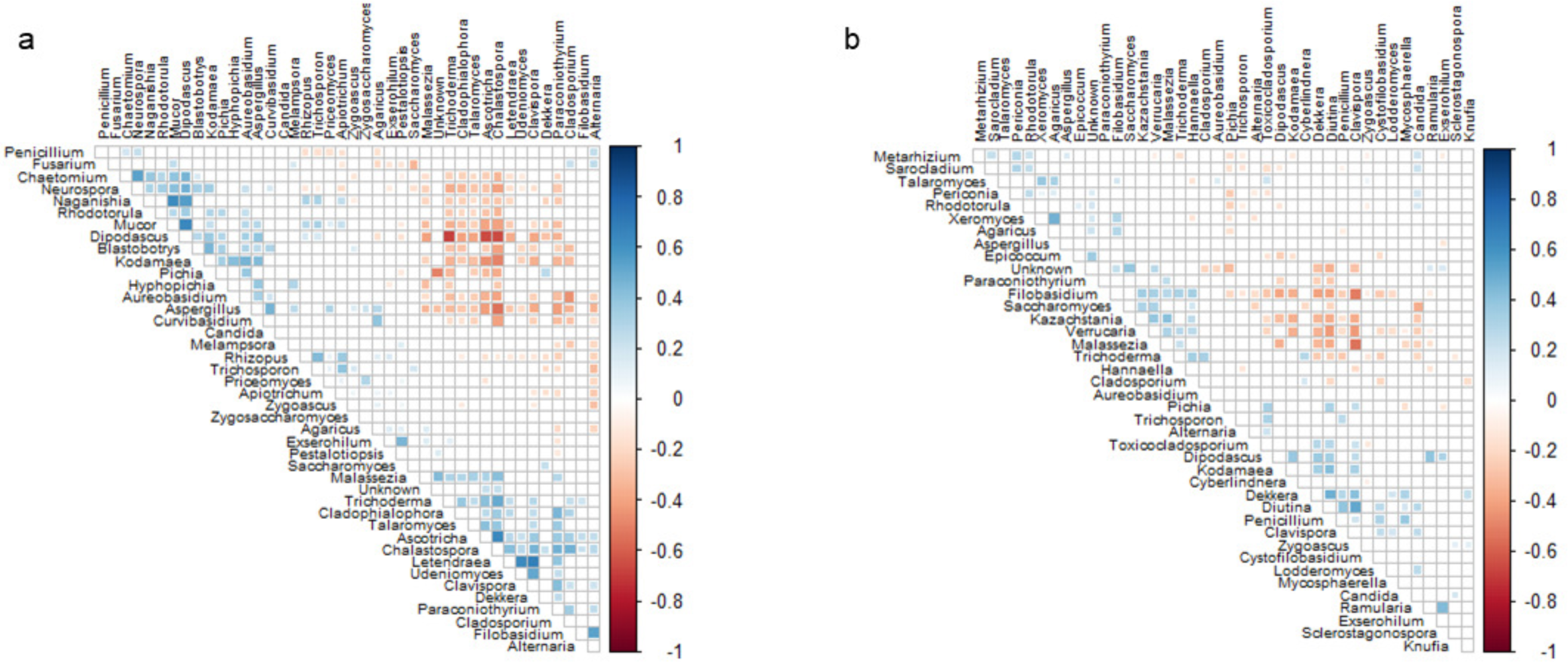
Correlations occurring between fungal taxa in **a**) rural and **b**) urban fungal mycobiota with P < 0.05 after FDR adjustment. Red squares represent significant negative correlations and blue squares represent significant positive correlations. The darker colours represent stronger correlations and non-significant correlations have been excluded from the plot.

### Biomarker taxa

Linear discriminant analysis (LDA) and the linear discriminant analysis effect size (LEfSe) (33) test for biomarkers was used to detect taxa that showed the strongest effect on group differentiation (**Figure 3a**). OTU level analysis uncovered 14 urban-associated species from 10 genera. Whereas, 17 rural-associated species from 11 genera, were detected as possible biomarkers. The most abundant rural-associated biomarker genera were *Hypopichia* and *Dipodascus*, with species *Hypopichia burtonii* and *Dipodascus geotrichum being the most ubundant* (**Figure 3b**). Whereas, the urban-associated biomarkers were dominated by the class *Tremellomycetes* and genera *Dekkera* and *Hannaella*. Whereas, species *Dekkera bruxellensis and Hannaella sinensis* dominated the urban-associated biomarkers.

## Discussion

The results from this study suggest that the gut mycobiome of the South African population is structured by geography and lifestyle. This finding is supported by the clustering of a large proportion of the fungal OTU’s into discrete rural and urban groups within the Venn diagram. Only a small percentage of OTUs were shared between the two populations, which may suggest that factors such as the environment, age and diet may play a role in shaping the differences in OTU clustering. These results were further corroborated by two distinct clusters, consistent with rural and urban locality.

Several studies have investigated the healthy human mycobiome (6, 7, 19, 25, 30, 34). In these studies, geography was not considered as a potential factor structuring the gut mycobiota. For instance, Nash et al. (2017) found no association between host phenotypic characteristics with mycobiome profile. This study suggests that diet, the environment, diurnal cycles, and host genetics may substantially influence the human gut mycobiome. However, the finding that the majority of the variation could not be explained by their metadata does suggest that other environmental factors, such as geography, may contribute to structuring the human microbiome (6).

Our study provides the first results showing the importance of geography in African populations. Geographic locality may be associated with different environmental factors, such as different climatic regimes, which may effect structural changes in the mycobiota. For example, climate significantly influences vegetation and farming practices and leads to region specific diets. These region-specific diets may ultimately influence the gut mycobiota. This is a reasonable prediction given previous findings showing that fungi have climate dependent biogeographic patterns (35, 36). These patterns are likely to determine the type of fungi individuals may be exposed to, which may in turn impact the colonization of fungi in the human gut. The most abundant rural-associated biomarker species found in this study, *Dipodascus geotrichum*, is ubiquitous in nature (37) whereas, *Hypopichia burtonii* is commonly isolated from corn, wheat, and rice (38). The urban-associated biomarkers were dominated by the species *Dekkera bruxellensis*, which are commonly isolated from fermented food such as wine, beer, feta cheese and sour dough (39-41). In contrast, *Hannaella sinensis* is commonly isolated from plants and soil (42, 43). The staple diet of the rural South African population primarily consists of a corn-based porridge (called ‘pap’). It is therefore not uncommon for a fungal species commonly isolated from corn to be a dominant biomarker for the rural population. Conversely, the urban population diet was more diverse and included fermented foods such as wine, sour dough bread and feta cheese, which are commonly available in supermarkets. Thus, the species *Dekkera bruxellensis* was identified as a dominant biomarker in the urban population.

In addition to geographic location, we found that smoking, mode of birth and breastfeeding significantly influenced gut fungal communities. Several studies have previously reported that these factors may significantly influence the initial colonization, subsequent composition and structure of bacterial members of the human gut microbiome (28, 44-46). Suhr et al. (2016) and Hallen-Adams et al. (2017) investigated the gut mycobiome of two cohorts that were exclusively on a vegetarian or a western diet. These studies found that the distribution of fungi differed considerably between the two cohorts (7, 47). Plant-associated *Fusarium, Malassezia, Penicillium* and *Aspergillus* species were detected at higher abundances within the vegetarian cohort, compared to the cohort on a conventional diet. The finding that smoking affected fungal community composition and structure is supported by several studies (48-50). The approximately 4000 chemical compounds produced by cigarettes have been shown to alter the composition of the gut microbiome (48, 50-53). The reported increase of *Clostridia* induced by smoking in murine models has also been indirectly confirmed in humans where an increased rate of *C. difficile* infection was greater in former and current smokers compared to never smokers (52). Moreover, the abundance of the fungus *Candida tropicalis* has also been reported to be significantly higher in *C. difficile* infection patients compared to healthy individuals. (54). The abundance of *C. tropicalis* has also been detected to be positively correlated with levels of anti-*Saccharomyces cerevisiae* antibodies (ASCA) (54). In our study *C. tropicalis* was detected to be higher in individuals who smoke compared to non-smokers whereas, the inverse was true for *S. cerevisiae.* These findings may confirm the antagonistic association between the species *C. tropicalis* and *S. cerevisiae*, as previously reported by Hoarau et al., (2016).

Most studies have identified the genera *Candida, Saccharomyces, Malassezia* and *Aspergillus* as the three most abundant in the gut of healthy individuals (6, 25, 32). To the best of our knowledge, our study is the first to report *Pichia* as one of the top four most abundant genera in the human gut mycobiome. This may be due to several factors including differences in cohort characteristics (e.g., geographical location, diet, genetic predisposition and climate). *Pichia* have been identified as both constituent members of the human oral (55, 56) and gut microbiome (34). Mukherjee detected a 1:1 abundance ratio in the oral mycobiome of individuals when *Candida* and *Pichia* were present together (56). Pichia was also observed to have an antagonistic effect against *Candida, Fusarium* and *Aspergillus*.

The yeast genera, *Pichia, Candida* and *Cladosporium*, dominated the South African gut mycobiome. Our findings agree with previous studies which show that members of the *Aspergillus, Candida, Debaryomyces, Malassezia, Penicillium, Pichia*, and *Saccharomyces* genera were the most recurrent and/or dominant fungal genera (34, 47, 57). In contrast to previous findings, our data indicate higher relative abundances of *Cladosporium*, detection of *Mucor* and the absence or low abundance of genera such as *Cyberlindnera*, and *Galactomyces* (6, 19, 58). Previous studies found that the gut mycobiome of a cohort from Houston, Texas, was dominated by *Saccharomyces, Malassezia* and *Candida* (6). By contrast, the genus *Malassezia* was not detected in the gut mycobiome of a Pennsylvania cohort, which was instead dominated by the genera *Saccharomyces* and *Candida* (19). Differences in study methodologies may be a source of these conflicting findings (6). One study amplified the Internal Transcribed Spacer 2 (ITS2) region of the fungal rRNA gene (6), and the second amplified the ITS1 region (19). Studies similar to the work by Gardes et al. (1993) and White et al. (1990), where ITS1F and ITS2 primer sets were used to amplify the ITS2 region, did not detect *Malassezia* (59, 60). The second reason for the observed differences has been attributed to differences in cohort characteristics, such as diet and/or geographical location. Strati (2016) and Raimondi’s (2019) investigating cohorts in Italy, detected same dominant fungal genera (58, 61), and the investigation of cohorts in two different states in the USA observed different results (6, 46). We used ITS1 and ITS4 in this study and found that the genera *Pichia, Candida* and *Cladosporium* dominated the urban cohort, whereas genera *Pichia, Candida* and *Aspergillus* dominated the rural cohort. The dominant taxa identified in urban and rural locations further support our assertion that geographic location plays a major role in the observed differences.

*Candida albicans* was the most dominant taxa n our cohort and is frequently reported as the most abundant *Candida* species in both diseased (62) and healthy individuals (63). *Candida* spp. not only colonize the gut (19, 34) but several other body sites, including the oral cavity (55, 64), vagina (65), and skin (66, 67). However, *Candida* are autochthonous to the mammalian digestive tract and species including *Candida albicans, C. tropicalis, C. parapsilosis*, and *C. glabrata* may grow and colonize at 37°C (7). A review of the literature suggest that *C. albicans* carriage in healthy individuals ranges from 30–60% (68) and that living mammals are considered a niche for these species as they are not found in significant concentrations in soil, food or air (69, 70). Raimondi et al., (2019) reported that *C. albicans* was frequently detected and dominated the cultivable mycobiota of different faecal samples (61).

## Conclusions

This study provides the first insight into the importance of geography and lifestyle factors on the gut mycobiome in rural and urban locations in Africa. We found that fungi in the gut display distinct patterns consistent with geographic locality. Redundancy analysis showed that several lifestyle factors were major drivers explaining the distinct community structure. The results of biomarker analysis revealed several ecologically important fungal taxa, which were unique to individuals from urban and rural areas. These results have significant health implications, particularly for immunocompromised individuals living in rural and urban locations. Such findings provide a valid basis for the development of novel therapeutics or preventative measures reliant on modulating the gut mycobiome.

## Methods

### Ethical clearance

All experiments were approved by the Ethics Approval Committee of the Faculty of Health Sciences at the University of Pretoria (EC 160630-051). Participants approved and provided informed consent prior to enrolment in this study. All experimental methods and experiments were in accordance with the Helsinki Declaration.

### Participant enrolment criteria for urban and rural areas

Volunteers were recruited from two rural locations and one urban location. For rural volunteers, we recruited individuals following traditional diets, with generally low levels of processed foods. Urban cohorts reported mixed diets and increased consumption of processed foods. Volunteers from the Ha-Ravele and Tshikombani villages located in the Vhembe District of the Limpopo Province comprised the rural cohort. Both villages are approximately 391 km and 439 km, respectively, from the closest city (Pretoria). This city, in the Gauteng province of South Africa, served as the urban sampling area (Figure 1a). In total, 100 stool samples were collected from healthy volunteers. These samples were equally divided between gender and locality [i.e. rural (25 males and 25 females) and urban (25 males and 25 females)]. Self-stool collection kits were provided to all volunteers (Easy Sampler® Stool collection Kit, Hounisen Lab Equipment A/S, Skanderborg, Denmark).

### Inclusion and exclusion criteria

The participants were all healthy adults age 18 – 50 years. Volunteers reporting antibiotic use/other treatments in the sample collection sheets were excluded from the study. Similarly, individuals who had been diagnosed with any inflammatory-related bowel diseases or gastrointestinal diseases within six months prior to sample collection were excluded from the study.

### DNA extraction

DNA was isolated using the PowerSoil DNA Isolation Kit (MO BIO Laboratories Inc., Carlsbad, CA) following the manufacturer’s specifications with minor modifications. Briefly, approximately 0.25g of stool sample was transferred into the Power-Bead tubes using a sterile disposable wooden spatula (Lasec Laboratories, RSA). The sample was homogenized by gently vortexing the tubes for 10 s before adding 60 µL of the lysis buffer. This was then incubated for 30 min. at 55°C prior to centrifugation at room temperature for 30 s at 10,000 × *g*. The supernatant from this step was transferred to sterile 2 mL tubes and 250 µL of inhibitor removal reagent was added to this. The samples were incubated on ice for 5 min., thereafter approximately 1.2 mL of binding buffer was added. Next, 70% ethanol (500 µL) was added and the contents precipitated by centrifugation at room temperature for 60 s at 10,000 × *g*. The DNA was eluted with 100 µL filter-sterilised autoclaved Millipore water and quantified using the NanoDro™ 2000/2000c Spectrophotometer (Thermo Scientific, Waltham, MA, USA). The quality of isolated DNA was confirmed by agarose gel electrophoresis, on 1% (w/v) agarose gel in 1 X TAE buffer (0.2% [w/v] Tris, 0.5% [v/v] acetic acid, 1% [v/v] 5 M EDTA [pH 8]) at 90 Volts for 45 min. in a BioRad Sub-Cell® GT gel electrophoresis system with gel red visualising agent. The gel was visualised using the BioRad Gel Doc system and viewed with a UV Trans-illuminator.

### ITS gene region amplification, sequencing and data processing

The internal transcribed spacer (ITS) region was amplified using fungal-specific primers (60): ITS1F (5’-CTTGGTCATTTAGAGGAAGTAA-3’) and ITS4 (5’-TCCTCCGCTTATTGATATGC-3’) and the HotStarTaq Plus Master Kit (Qiagen, Valencia, CA). Amplicons from different samples were pooled to equimolar concentrations and purified of short fragments using Agencourt Ampure beads (Agencourt Bioscience Corporation, USA). Paired-end 2 × 250bp sequencing was performed on an Illumina MiSeq instrument (Illumina Inc., San Diego, CA, USA) at Mr DNA (Shallowater, TX 79363).

The resultant data were analysed using the Quantitative Insights into Microbial Ecology (QIIME2) software version 2018.8.0 (71). Demultiplexed sequences were assessed for quality and those shorter than 200 bp, with quality scores below 25, containing more than two ambiguous characters or more than one mismatch to the sample-specific barcode or the primer sequences, were excluded from further downstream analyses. Chimeric sequence detection and operation taxonomic unit (OTU) selection was performed at 97% sequence similarity using USEARCH v11 (72). Taxonomies were assigned to each OTU using the UNITE (release 7_99) databases for fungi (73). Singletons were excluded, and each sample was randomly subsampled (rarefied) to the same number of sequences per sample (17 980).

### Statistical analyses

All statistical analyses were performed in R version 3.5.1 using R studio (74, 75). Alpha diversity indices (observed, Chao1, Shannon, and Simpson indices), together with rarefaction curves were calculated and visualized using the R packages “phyloseq” and “ggplot”. First, the Shapiro-Wilk’s test was applied to determine data distribution (76). Subsequently, the unpaired two-sample Wilcoxon rank sum test (77, 78) was applied to determine significant differences between the alpha diversity indices using the R packages “dplyr” version 0.4.3 and the “ggpubr” version 0.1.8. (79, 80). In these analyses, the rural or urban location was specified as a random factor.

The R packages “phyloseq” (81) and “microbiomeseq” (82) were used to calculate and visualize relative taxa abundance at phylum and class level. OTU abundance was transformed to relative abundance and taxa with relative abundance less than 0.1% were removed. The Wilcoxon rank sum test was applied to determine significant differences between taxa in the urban and rural samples. Whereas, the Kruskal-Wallis test (83) was applied to determine significant differences between taxa in the four sample types.

The local contribution to beta diversity (LCBD) was calculated according to (84). The LCBD describes the degree of uniqueness of a given sample in relation to the overall community composition. The taxa abundance was normalized to obtain the proportion of most abundant taxa per sample. Location was used as the grouping variable and the Hellinger method was used for the dissimilarity coefficients calculation.

Pairwise similarities among samples were calculated using the Bray–Curtis index of similarity. The resulting matrix was represented visually in a nonmetric multidimensional scaling (NMDS) plot to observe community structure. Using the vegan package (85), a permutational multivariate analysis of variance (PERMANOVA) (86) based on 9999 permutations of the data, was performed to test whether differences between sample groupings in the NMDS ordinations were statistically significant. Microbial community similarities and the homogeneity of dispersion between the rural and urban sample groups were tested using the ANOSIM and ADONIS tests, respectively (87, 88).

The effect of the different recorded environmental factors on fungal community composition and structure was determined through redundancy analysis (RDA). First, the OTU-count data were Hellinger-transformed. The contribution of highly correlating OTUs (*p*-Value < 0.05) with redundancy axes was identified using the envfit function from the R package vegan (85). To measure the relationship of abundant taxa with measured anthropometric factors (age, BMI, height, and weight), a Spearman correlation analysis was done and visualized in the R package microbiomeseq (82).

Fungal-fungal relationships were interrogated using SparCC (89). Correlation was based on measuring the linear relationship between log transformed abundances. First, data were filtered to remove OTUs that had less than 2 reads on average. SparCC was used to generate true correlation coefficients from which pseudo p-values were calculated. The calculate pseudo p-values were false discovery rate (FDR) adjusted (90) and the correlation matrix was visualized using the “corrplot” function (91) in R.

Potential biomarker taxa which differed in abundance and occurrence between the two geographic groups were detected by linear discriminant analysis (LDA) effect size (LEfSe) (33). The LEfSe was calculated using the online Galaxy web application (92) with the Huttenhower lab’s tool (https://galaxyproject.org/learn/visualization/custom/lefse/). First the nonparametric factorial Kruskal–Wallis sum rank tests (alpha = 0.01) was used to detect differential abundant features (at genera, family, class and phylum level) within the two geographic locations (rural and urban). The phylogenetic consistency was then tested using the pairwise Wilcoxon rank-sum tests (alpha = 0.01). Finally, the effect size of each differentially abundant feature was estimated using the LDA. The all-against-all classes were compared (most stringent) and a linear discriminant analysis score value of 2.0 was chosen as threshold for discriminative features.

### Consent for publication

All participants provided consent for publication of study results of the collected biomaterials paired with anonymized information on age, sex, location, diet and other data.

## Availability of data and materials

The sequence data generated in this study are available on the NCBI (https://www.ncbi.nlm.nih.gov/) under the following accession number: PRJNA589500.

## Competing interests

The authors declare that they have no competing interests.

## Acknowledgements

This research was supported by the South African Medical Research Foundation (TPM). MHK and KM received scholarships from the National Research Foundation. We wish to thank the Fulbright program for providing sabbatical support to TPM, JJ was funded by the Microbiomes in Transition (MinT) initiative at the Pacific Northwest National Laboratory in Richland, WA, USA. PNNL is operated for the DOE by Battelle Memorial Institute under Contract DE-AC05-76RL01830. We would like to thank all the volunteers for donating their data, time and samples to our research.

## Supplemental Figure legend

**Figure S1** Comparison of mycobiota between urban and rural participants a) Venn diagram showing the unique and shared phylotypes for samples collected from urban and rural participants. b) Rarefaction plot showing sequencing coverage. c) Taxa abundance data was normalised to obtain the proportion of most abundant taxa per sample. The diameter of the points at the bottom of the plot corresponds to the magnitude of the LCBD value for a particular sample. The bars correspond to taxa that are most abundant with the top taxa sharing a bigger portion of the bar for each sample.

